# Heart Scar-In-A-Dish: Tissue Culture Platform to Study Myocardial Infarct Healing In Vitro

**DOI:** 10.1101/2025.02.28.640625

**Authors:** MJ Potter, JD Heywood, SJ Coeyman, WJ Richardson

## Abstract

Myocardial Infarction (MI) is a major contributor to morbidity and mortality, wherein blood flow is blocked to a portion of the left ventricle and leads to myocardial necrosis and scar formation. Cardiac remodeling in response to MI is a major determinant of patient prognosis, so many therapies are under development to improve infarct healing. Part of this development involves in vitro therapy screening which can be accelerated by engineered heart tissues (EHTs). Unfortunately, EHTs often over-simplify the infarcted tissue microarchitecture by neglecting spatial variation found in infarcted ventricles. MI results in a spatially heterogeneous environment with an infarct zone composed mostly of extracellular matrix (ECM) and cardiac fibroblasts, contrasted with a remote (non-infarct) zone composed mostly of cardiomyocytes, and a border zone transitioning in between. The heterogeneous structure is accompanied by heterogeneous mechanics where the passive infarct zone is cyclically stretched under tension as the remote zone cyclically contracts with every heartbeat.

We present an in vitro 3-dimensional tissue culture platform focused on mimicking the heterogeneous mechanical environment of post-infarct myocardium. Herein, EHTs were subjected to a cryowound injury to induce localized cell death in a central portion of beating tissues composed of neonatal rat cardiomyocytes and cardiac fibroblasts. After injury, the remote zone continued to contract (i.e., negative strains) while the wounded zone was cyclically stretched (i.e., positive tensile strains) with intermediate strains in the border zone. We also observed increased tissue stiffnesses in the wounded zone and border zone following injury, while the remote zone did not show the same stiffening.

Collectively, this work establishes a novel in vitro platform for characterizing myocardial wound remodeling with both spatial and temporal resolution, contributing to a deeper understanding of MI and offering insights for potential therapeutic approaches.

## 1. Introduction

Heart disease is the primary cause of death in the United States, accounting for 696,962 deaths in 2020 (20.6% of total) and $239.9 billion in healthcare costs and lost productivity.^1,2^ Myocardial infarction (MI) is a significant contributor to the prevalence and burden of heart disease. MI impacts more than 805,000 Americans annually and occurs when coronary blood flow becomes obstructed, leading to ischemic damage in the downstream myocardial tissue.^2^ Following MI, damaged cardiomyocytes undergo degradation and removal, while cardiac fibroblasts infiltrate the infarct to deposit extracellular matrix proteins essential for wound repair and scar formation.^3–8^ This process stabilizes the wound area to endure forces during the cardiac cycle but creates tissue that is mechanically, electrically, and biologically distinct from healthy myocardium.^9–11^ Consequently, understanding the tissue response to MI is critical for developing effective treatments. There is a clear need to characterize these heterogeneous changes in myocardial tissue.

Traditional 2D culture techniques offer cost-efficient, high-throughput systems that are simple to observe and measure. However, these systems cannot facilitate the full extent of tissue complexity, as they limit cell-cell and cell-environment interactions to a single plane.^12–14^ In contrast, 3D culture techniques provide numerous benefits including higher capacity for morphological and physiological development, scaffold designs that better support cellular differentiation and maturation, and increased structural complexity for a more physiologically accurate environment.^12–15^ This complexity is particularly advantageous for cardiac tissues, which exhibit diverse environmental characteristics. These characteristics include mechanical, electrical, and biological properties which are detected by embedded cardiomyocytes and cardiac fibroblasts. Mechanical forces applied in vitro serve to align cells, enhance myocyte contraction, and promote fibroblast differentiation.^16–24^ Simulated MI wounding triggers localized necrosis and apoptosis, myofibroblast activation, and ECM turnover.^25–32^ Various treatments and stimulations such as selective ablation for atrial fibrillation, cardiomyocyte maturation through electrical conditioning, and handling of calcium signaling are also possible in 3D EHT based experiments.^33–36^

Given the heart’s mechanical activity, greater comprehensive mechanical characterization of EHT systems can improve study relevance and reveal further insights into the impacts local mechanics have on post-MI processes. Although bulk force changes during systole are expected, distinct regions of healthy, infarcted, and border tissue mechanics should be characterized by a precise spatial distribution of forces. Additionally, to further enable the study of dynamic mechanics of a full contraction cycle, EHT systems can benefit from externally controlled pacing which can provide human-relevant cardiac cycle timings. This electrical stimulation can prove more adjustable and consistent than attempts to implement native pacemaker cells.

To address these challenges, we propose a 3D EHT MI platform incorporating both spatial and temporal analysis for enhanced biomechanical accuracy. Motivated by cryo-injury methods previously developed for in vivo and in vitro wound models, we induced cryo-wounding in our EHTs to generate three distinct regions, or zones: wound, remote, and border.^37,38^ These different zones possess unique properties arising from cell distributions, which yield distinct force generation and systolic strains throughout the cardiac cycle. Over 7 days of culture, the tissue microenvironments evolve toward biomechanical homeostasis, providing a first-step platform capable of analyzing long-term ECM remodeling across different mechanical regions and the resultant heterogeneous mechanical properties such as force generation, stiffness, and deformation.

## 2. Methods

### A. Platform Assembly

Custom 32-well plates, adapted from a previously published platform, were printed with a Formlabs Form2 3D printer (CAT# PKG-F2) using Dental LT V2 biocompatible resin (CAT# RS-F2-DLCL-02). Upon completion of manufacturer recommended post-processing, a thin layer of the same resin was applied to the bottom surface of the printed plate. This was used to affix the plate to 10cm square petri dish. The adhered plate was then cured at the manufacturer recommended temperature and duration under UV light within the Form Cure (CAT#FH-CU-01, Formlabs, USA).

Free-floating pistons were similarly printed and cured. A 1mm cube magnet (SuperMagnetMan CAT# C0010) was placed into a slot on top of each piston and filled with a small amount of Dental LT V2 resin combined with contrast dye. Pistons were cured under UV at room temperature instead of high heat to avoid demagnetizing the embedded permanent magnets. Following an initial 30-minute cure, a second layer of resin, without contrast dye, was placed over the initial layer and cured for another 30 mins.

Prior to use, all parts underwent a cleaning and sterilization protocol consisting of sequential washes for 15 minutes of 10% bleach, detergent, and 70% ethanol. Each device was assembled in a biosafety cabinet. Carbon electrodes were placed into grooves found at opposing sides of the plate and anchored in place using ½” stainless steel screws. One screw also secured a conductive ring terminal to the carbon electrode block. The electric pulse generator was connected to the ring terminals of each electrode by 16ga stranded copper wire. Parafilm was used to seal the gap around each wire to preserve sterility. Then spacers were placed into each of the 32 culture wells. These spacers acted as a backstop upon which the pistons would rest, resulting in a predetermined gel length that left space behind the piston within the culture well. This space would allow stretching of samples during the culture process. Pistons were placed into each well of the 32-well plate with the back of each piston in direct contact with the inserted spacers. After the assembled platform was placed in a magnetic alignment tray which pulled the pistons against their spacers, each well was then filled with 100 µL of 3.5% Pluronic F-127 in PBS for 30 mins before removing the solution. The platform was allowed to dry and underwent a final 15-minute UV sterilization step. Figure 1(a) displays the assembled platform.^39^

**Figure 1.**
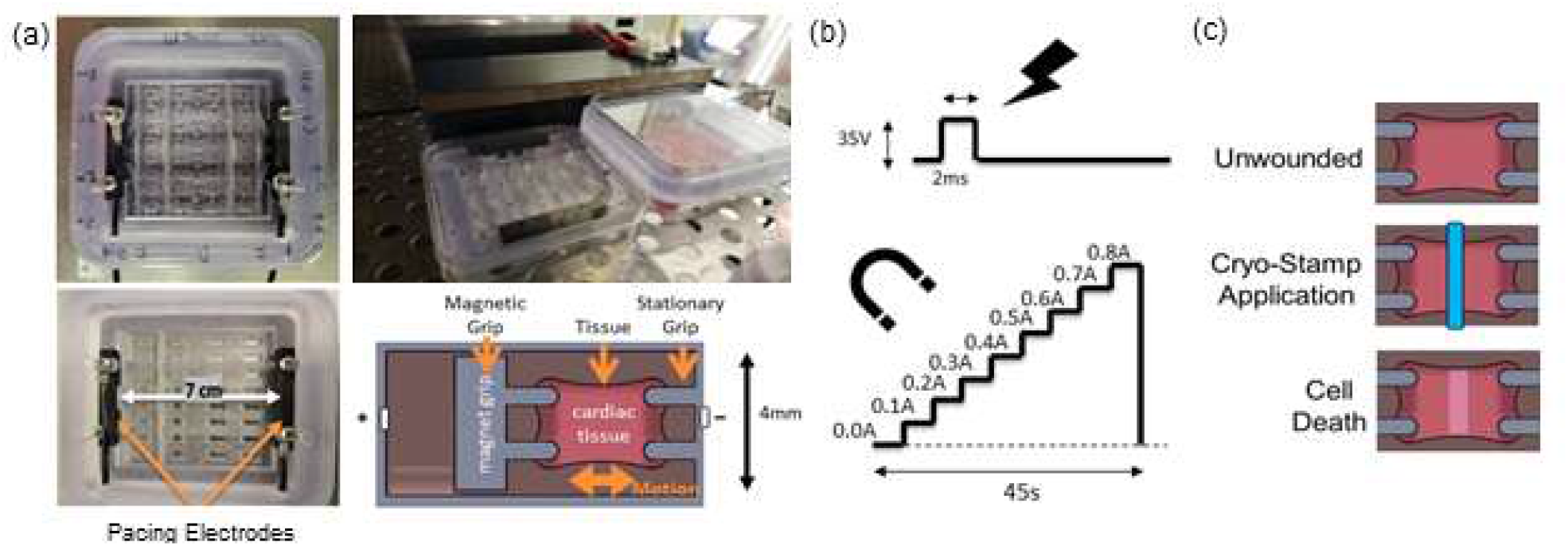
Culture Platform and schematic representation of stimulation modalities. (a) Top-down view of the culture platform showing the well arrangement and carbon electrode positioning. (b) Illustration depicting the 2 ms unipolar 35V electrical stimulation pulse and the electromagnet discrete amperage ramp program utilized to stretch the engineered heart tissue constructs. (c) Illustration of the cryo-wounding process.

### B. Hydrogel Fabrication and Seeding Parameters

Primary neonatal rat cardiomyocytes and cardiac fibroblasts were sourced from the ventricles of neonatal Sprague-Dawley rats (Jackson Labs, USA) birthed 1-3 days prior to harvest as approved by the Institutional Animal Care and Use Committee at Clemson University. Ventricular segments were rinsed in HBSS on ice, transferred to a trypsin solution for mincing, and the resultant homogenate was digested overnight at 4°C in fresh trypsin solution. Following the overnight digestion, 5 mL of NCIS Purified Collagenase Solution in L-15 media was added and the solution was agitated for 40 minutes at 37°C. Digested tissue was then filtered using a 70um strainer, and resulting cells were incubated at room temperature, loosely capped in a biosafety cabinet, for 40 mins. Cells were then pelleted at 200g for 5 mins. After resuspending the pellet in 40mL of DMEM + 10% FBS + AB/AM, the cell solution was equally distributed among four 10cm petri dishes and incubated at 37°C for 2h.

After incubation, the supernatant was pipetted over the solution 10 times to remove any loose cells. The supernatant (now cardiomyocyte-enriched portion) was transferred to a 50mL conical and counted using a hemocytometer. Adherent cells (fibroblast-enriched portion) were allowed to continue in culture and were passaged after 3 days. Harvested cardiomyocytes were utilized immediately while the cardiac fibroblasts were utilized at passage 2 arising from the preceding harvest.

Cells were counted using a manual hemocytometer. Each EHT was fabricated with 525K cardiomyocytes (P1) and 175K cardiac fibroblasts (P2). Cells were resuspended in 0.44U/mL thrombin. Then, an equal volume of 5.5mg/mL fibrinogen in HEPES-BSS solute was added to cell-containing thrombin solution. Immediately, 85uL of the EHT solution was pipetted into each well of the 32-well plate. The EHTs were polymerized for 60 mins at 37°C before adding gel media (DMEM, 10% horse serum, 33 µg/ml aprotinin, 50µg/ml L-ascorbic acid, 0.1% insulin, 1% pen-strep, 0.2%ampho-b) followed by overnight incubation. After one day, the piston spacers were removed and media was replaced. Media was changed every second day throughout the time course.

### C. Cryowound Injury

A custom injury tool was printed using the Formlabs Form2 printer and Dental LT V2 resin. The injury tool contained 1mm-wide segments that bisected each EHT at its midpoint. To apply the injury, media was first aspirated from the plate. The injury tool was then submerged in liquid nitrogen for 60 seconds. It was then removed and held against the EHTs for 60 seconds. The media was replaced following wounding.

### D. Electrical and Mechanical Stimulation

On the third day following EHT fabrication, the media was changed, and the magnetic alignment tray was removed. Using an IonOptix Myopacer, a 1Hz, 35V, 2ms duration positive pulse waveform was applied to the electrodes, shown in figure 1(b). This voltage was used to supply 5V for every centimeter of separation that existed between the electrodes in culture. Once pacing was initiated, a McMaster-Carr electromagnet (EM) (CAT# 5684K23) was placed 3mm away from the outer edge of the culture plate supplying a constant force supplied by a constant 0.3A from a computer-controlled DC power supply (Siglent CAT# SPD3303X-E). Calibration of forces applied to the embedded permanent magnets by the electromagnet have been described in previous work.^40^

### E. Immunofluorescence Staining

For IF staining, media was removed from the plate and 16mL of 4% paraformaldehyde (PFA) was added for overnight fixation. After next day PFA removal, tissues were washed 3x with PBS and permeabilized with 0.2% Triton-X100 for 30 mins at room temperature. After 3 further PBS washes, tissues were blocked using Pierce Blocking Buffer (PBB) overnight at 4°C. The blocking buffer was then removed, tissues were washed 3x with PBS, and a primary antibody solution was added containing fibroblast specific protein (FSP-1) (1:150) and cardiac troponin (CT-1) (1:200) in PBB. Plates were then covered and incubated overnight at 4°C. After washing 3x with PBS, a secondary antibody solution was added containing Goat-anti-Rabbit Alexa Fluor 594 (1:1000) and Goat anti-Mouse Alexa Fluor 488 (1:2000) in PBB. Tissues were covered and incubated overnight at 4°C. After washing 3x in PBS, a DAPI stain (1:47,667) in PBB was added for 20 mins, while rocking at room temperature. After washing 3x with PBS, tissues were imaged at 4x magnification through Texas Red, Green Fluorescent Protein, and DAPI filter cubes with an EVOS FL Auto microscope (Life Technologies Carlsbad, CA).

### F. Mechanical Evaluation

An EVOS FL Auto microscope with accompanying PearlScope64 software was used to record videos of each EHT. Using a custom imaging harness, each 32-well plate was placed onto the microscope stage with the Myopacer leads still attached with an applied 1 Hz pacing cadence. A discrete increase in amperage levels from 0.0A to 0.8A in 0.1A increments every 5 seconds, shown in figure 1(b), was supplied to the EM, which was again placed 3mm from the base of the plate. This discrete amperage ramp was controlled through custom LabVIEW (NI Base) software and the tissue deformation resulting from the applied tension was recorded. Videos were taken using the 2x objective, and the files were created and saved through a screen-capture software at a resolution of 1920x1080 and a framerate of 60fps. Videos were taken of each gel on each plate and imaged on the following timepoints: Prewound (0d), 1d, 3d, and 7d.

ImageJ was used to measure the baseline unstretched gel widths at the middle and end positions. Discrete widths along the gels were interpolated for the positions falling between the middle and end positions via a constructed linear relationship between the central and terminal points. The interpolated widths of the gels were used to calculate a corresponding compaction factor by determining the ratio of interpolated width to the original gel width of 4mm. Gel thicknesses were then estimated by applying each position’s compaction factor to the original gel thickness, determined by the height of the culture well, 3.25mm. The cross-sectional areas were calculated using these thickness and width estimates.

For systolic strain analysis, individual frames of ‘systole’ and ‘diastole’ were manually chosen. These frames were then extracted at native resolution and imported into NCORR 2D digital image correlation MATLAB software.^41^ The median Lagrangian surface-mapped strain value (along the primary long-axis parallel to bulk tissue motion) was found from across the 0.5mm central portion of the short-axis for discrete datapoints along the long-axis. This was performed along the central 90% of the long-axis to avoid edge artifacts that could arise from the mechanical restraints at each respective end of the gel construct.

Stress values were calculated by dividing the experimentally determined force values by the estimated cross-sectional area for each discrete position along the long-axis of the gel construct. Stresses and strains were averaged within seven bins of uniform width, and a linear regression was performed to attain the estimated modulus of elasticity value for each individual region within each sample.

### G. Statistical Analysis

We tested a total of 60 tissue constructs divided into six experimental groups: wounded and unwounded control cultured for 1-day, 3-day, or 7-days. To test for spatial differences in cell density, systolic strains, and tissue stiffnesses within the gels, a repeated measures ANOVA was applied to the three distinct regions (wound zone vs. border zone vs remote zone) within each experimental group, followed by post-hoc paired t-tests to find statistically significant differences between regions (p-value < 0.05).

To evaluate the effects of wounding on cell density, fluorescent intensity of cell markers at 1-day, 3-day, and 7-day timepoints was normalized as a fold-change relative to each respective region’s 0-day intensity (pre-wound). A one-sample t-test against a hypothetical mean of 0 was then run for each region within each experimental group to find statistically significant changes in cell density after wounding (p-value < 0.05). Lastly, to test for systolic strains during tissue beats, a one-sample t-test against a hypothetical mean of 0 was run for each region within each experimental group (p-value < 0.05).

## 3. Results

### A. Immunofluorescent Staining and Cellular Density of Wounded Tissues

Control samples showed a mostly uniform distribution of cells across all regions in each tissue. Slight increases in both cardiomyocyte and cardiac fibroblast signal intensity across all regions between day 0 and day 7 (figure 2 (b) and (c)) were observed. In contrast, wounded tissues showed significant reduction in cardiomyocyte and fibroblast signal across all spatial regions at day 1 and day 7 post-wounding. The degree of reduction was spatially heterogeneous across the wound, border, and remote regions (figure 2 (a), (b), and (d)). Notably, both cell type concentrations in the wounded region were reduced by ∼40% after 1 day. Concentrations were reduced by ∼30% in the border zone and by only ∼5-10% in the remote zone. This pattern was mostly maintained through day 7 with the exception that fibroblast levels in the wound and border zone slightly recovered between day 1 to 7. However, cardiomyocyte levels continued to drop between day 1 to 7.

**Figure 2.**
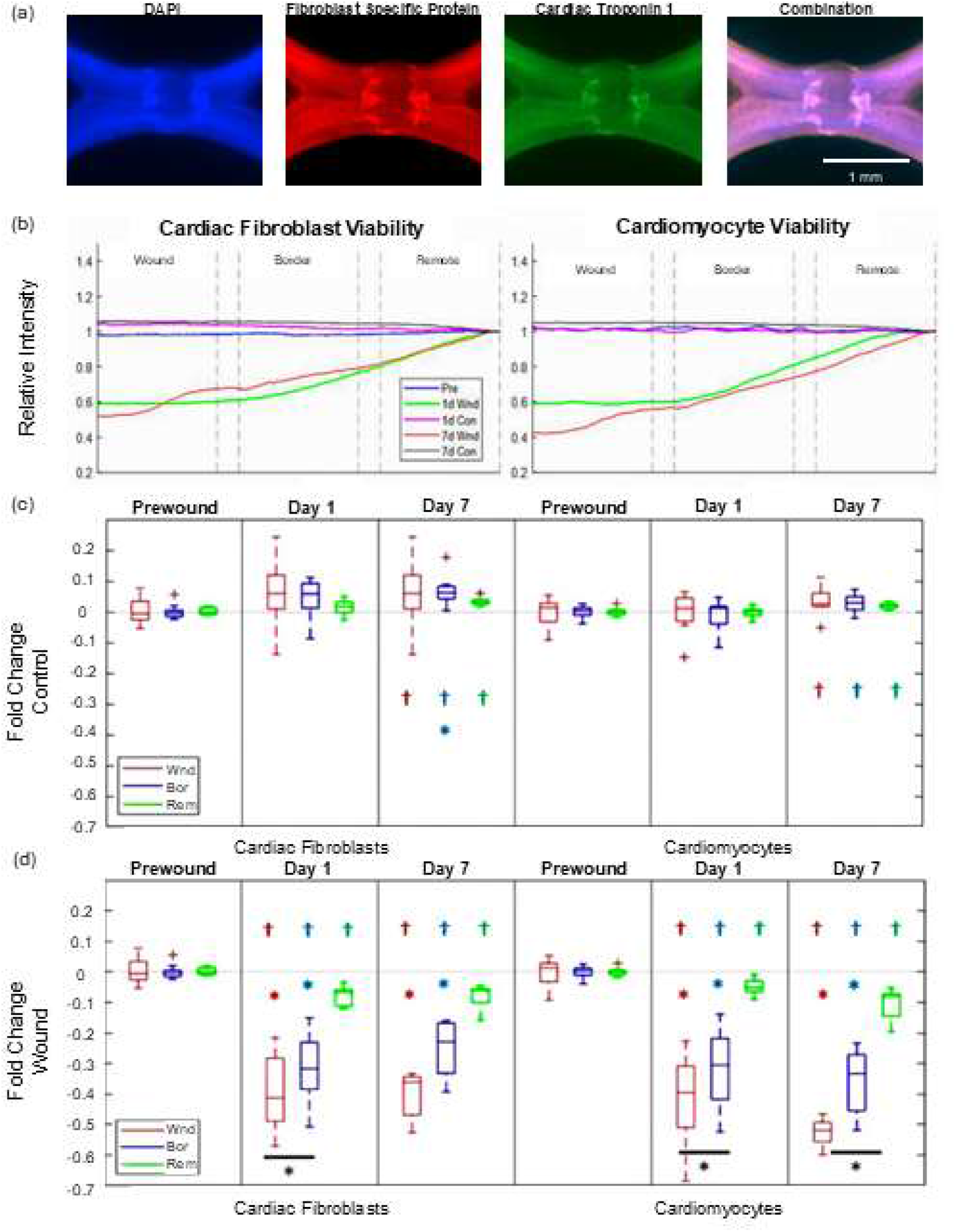
Cryowound Injury of Fibroblast and Myocyte Populations Across Different Regions. (a) Representative immunofluorescence staining of DAPI (nuclei), FSP-1 (fibroblasts), cTn-1 (cardiomyocytes), and the combined overlay of a wounded tissue after 7 days of culture. (b) Quantification of FSP-1 and cTn-1 fluorescence indicating cell concentrations across the wound, border, and remote zones. Each sample has been normalized to a 10 pixel-wide subset at its own outer edge. For statistical comparisons, the Area Under the Curve (AUC) from these map traces ((b) curves) was quantified for each region of each sample, normalized by each individual sample’s prewound AUC, and compared for control samples (c) and wounded samples (d). * indicates significant difference from remote zone (p < 0.05), † indicates significant difference from hypothetical mean of zero (p < 0.05), and * with bar indicates significant difference between wound and border zones (p < 0.05). Sample sizes were: prewound (n = 60), 1d Control (n = 36), 7d Control (n = 9), 1d Wound (n = 24), 7d Wound (n = 9).

### A. Systolic Strain Analysis

Surface deformation mapping captured strains between peak systolic contraction and diastolic relaxation. These strains were used to evaluate the effects wounding had not only on passive EHT mechanics, but also on how active contractility mechanics were impacted. As shown in figure 3, EHTs exhibited contractile strain throughout the 7-day culture period in unwounded control samples.

**Figure 3.**
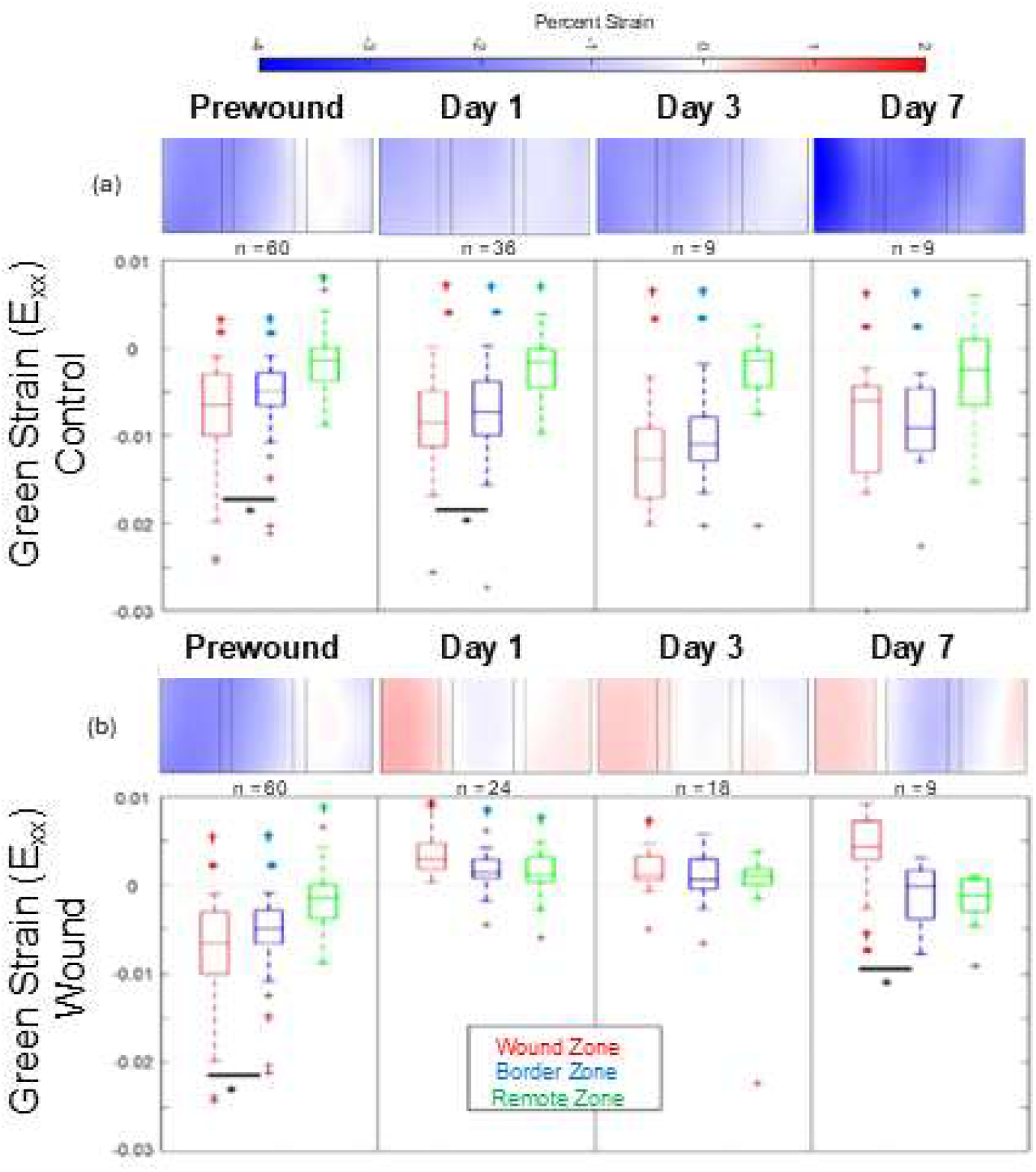
Systolic Strain Analysis. (a) Control samples (top row) exhibited contractile strains throughout the tissue, with the most significant contraction seen in the center of the EHT. (b) In contrast, the wounded regions of the cryoinjury samples showed tensile strains indicating the wounds were being cyclically stretched each time the remote zones continued to contract. * indicates significant difference from remote zone (p < 0.05), † indicates significant difference from hypothetical mean of zero (p < 0.05), and * with bar indicates significant difference between wound and border zones (p < 0.05). Sample sizes were: prewound (n = 60), 1d Control (n = 36), 3d Control (n = 9), 7d Control (n = 9), 1d Wound (n = 24), 3d Wound (n = 18), 7d Wound (n = 9).

In contrast, wounded samples exhibited stark heterogeneity with average strains in the wound region shifting from contractile (negative) pre-wound to tensile (positive) post-wounding at day 1. These tensile strains became significant by day 7 while the remote zone continued to demonstrate contractile strains even in the wounded tissues.

### B. Stiffness Analysis

Regional stiffnesses displayed in figure 4(b) begin fairly uniform with no statistical differences between values across subregions or conditions observed within the prewound samples. This changes for observations made on day 1 following wounding. While the lack of variation is maintained for control samples on day 1, the infarct subregion of the wounded samples was observed to be stiffer than either of the other analyzed regions within the gel. The wounded gels continued on a stiffening trend that resembled a stairstep descending from wound to border to remote. Repeated measures ANOVA confirmed that stiffnesses were significantly dependent on region across the week-long time course for the wounded tissues (p<0.05), but stiffnesses were not significantly dependent on region in the unwounded tissues.

**Figure 4.**
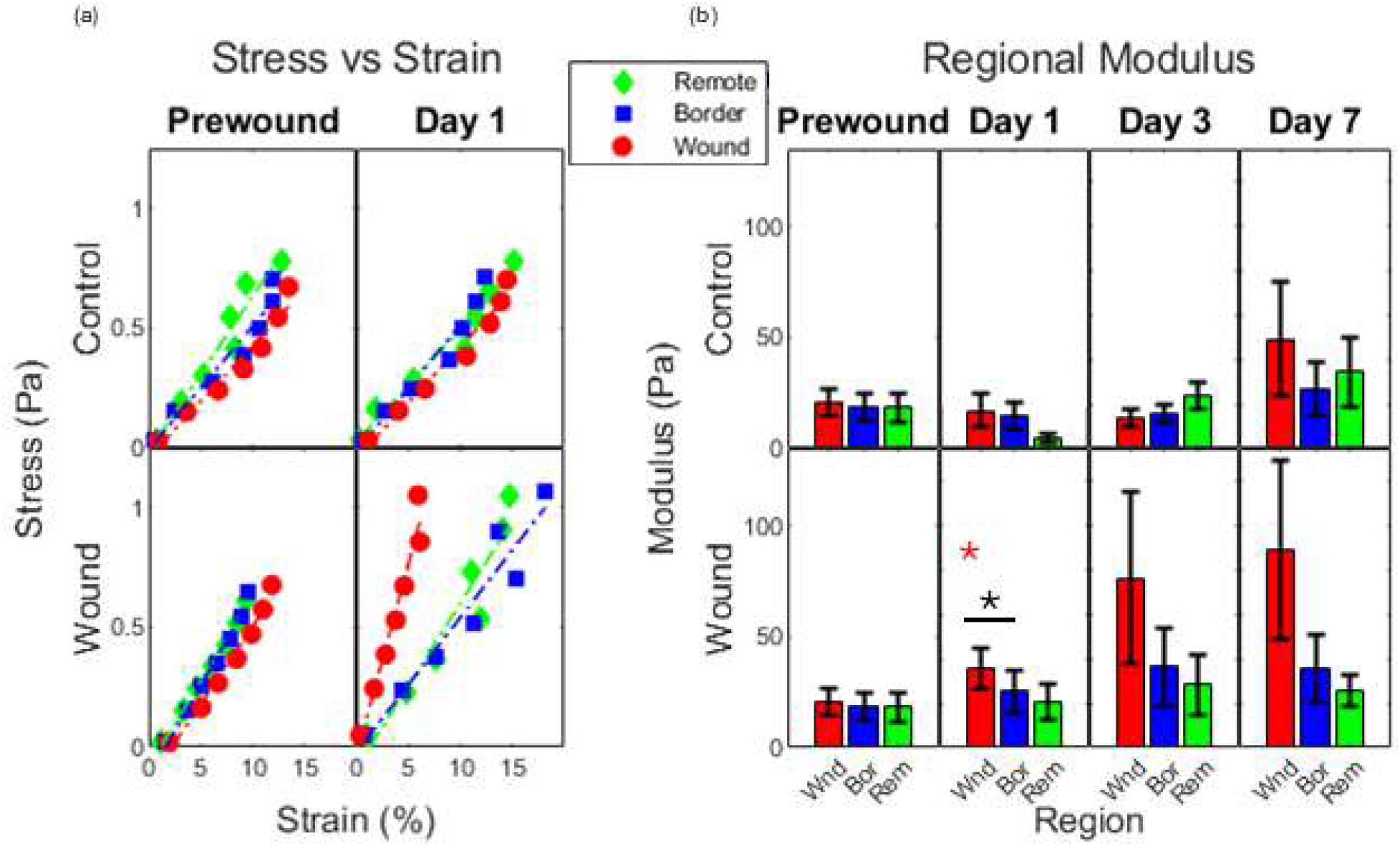
Regional Mechanics. Stiffness relationships from the subregions dividing the central 2.3mm sample length. Analysis sub-regions are 0.3mm in length and separated by gap-regions that are 0.2mm in length. (a) Representative stress-strain plots from individual samples are presented. B) Modulus values for each region were calculated by linear regression for stress-strain values of individual samples. * indicates significant difference from remote zone (p < 0.05) and * with bar indicates significant difference between wound and border zones (p < 0.05). (Wounded n = 6 | Control n = 4).

## 4. Discussion

This paper presents an in vitro heart attack model using a cryowound technique to induce cellular death in a 3-dimensional EHT. This results in a spatially heterogeneous tissue construct possessing wound, border, and remote zones with variation in cell density, tissue strains, and tissue stiffnesses across these regions. The healing infarct in vivo is without question a very complicated environment with dynamic mechanics, varied geometries, and spatial heterogeneity. This complexity has confounded therapeutic strategies for improving cardiac remodeling post-MI. Even carefully controlled experimental animal infarct models vary wildly in collagenous scar structure, mechanics, and chronic remodeling. This translates to difficulty ascertaining possible therapeutic efficacy without testing across a range of species, infarct locations, and reperfusion protocols.^11^ This reality suggests that a simpler, more consistent, and more widely applicable methodology of evaluating therapeutic efficacy is needed to help elucidate the biology of the infarcted heart and to help accelerate therapeutic strategies which improve cardiac function post-MI. The experimental platform presented within this study is a preliminary investigation into the natural progression of a simplified engineered cardiac tissue construct following myocardial injury. As literature findings clearly demonstrate the importance of wall mechanics, investigating the progression of the regional deformations and stiffnesses found within our mechanically active constructs is an important contribution towards an improved understanding of early infarct mechanics progression.

The normal, healthy myocardium is composed primarily of cardiomyocytes and cardiac fibroblasts. The production of mechanical contractile forces to pump blood out of the heart during systole is the primary function of cardiomyocytes. Maintenance of the extracellular matrix that serves as a structural scaffold for cells to organize around is the primary function of fibroblasts. Importantly, both cell types translate mechanical stimuli into biochemical signals and are highly sensitive to their mechanical environment. Mechanical stimuli trigger changes in cell morphology (cell size, myofilament length, sarcomeric actin), gene expression, and functional activity.^42–44^ Environmental forces directly influence fibroblast proliferation and differentiation, ECM synthesis, and matrix remodeling through mechanoreceptors such as integrins and cadherins, which are crucial components of cellular adhesion and communication pathways. The mechanical forces from cardiomyocytes can inform fibroblast behavior, affecting ECM composition and mechanics. Conversely, the remodeled ECM affects cardiomyocyte loading conditions, force generation, and electrical activity. This homeostatic balance within the myocardium reflects the interplay of mechanical and biochemical signaling required to sustain healthy tissue. It also points towards the underlying pathophysiological mechanisms by which cardiac remodeling and function can change due to cardiovascular disease or therapeutic intervention.

Following infarction, the dynamic architectural and functional attributes of the myocardial tissue undergo a significant transformation. Tissue ischemia leads to cellular necrosis in the infarct zone driven by hypoxia, oxidative stress, and inflammation.^45–47^ Fibroblasts are recruited to the site of injury via chemotaxis, progenitor cells, differentiation, and other cellular responses. ^48–52^ These cells, typically quiescent within the healthy myocardial matrix, activate as myofibroblasts and are characterized by enhanced proliferation, migration, and production of extracellular matrix proteins. Myofibroblasts are crucial for producing and depositing these extracellular matrix proteins, providing structural integrity and stability to the damaged myocardial tissue.

Myofibroblasts regulate the balance of ECM deposition and degradation (for example, MMPs and TIMPs which can exhibit varied activity based on applied mechanical strains), ensuring the formation of a wound-stabilizing fibrotic scar.^53^ For this, they secrete several matrix proteins (predominantly collagen, but also proteins such as elastin, fibronectin, and vitronectin) which form a structural scaffold reinforcing the infarcted tissue.^54–57^ This fibrotic response, while essential for preventing rupture and maintaining the structural integrity of the myocardial wall, often culminates in tissue stiffening. Importantly, this remodeling response results in heterogeneous mechanical environments. While the remote, uninfarcted, cardiomyocyte-rich region continues to contract with each heartbeat, the infarcted zone is now a passive, scar-rich region that is stretched by the remote contractions.^9,10^ The distinct tissue regions - healthy, infarcted, and border zone - are characterized by unique mechanical and biological properties and thus should be considered separately when discussing post-infarct wound remodeling.

While previous work has tested the regional effects of hypoxia on cardiac tissue in vitro, we sought herein to develop an in vitro EHT platform that could recapitulate heterogeneous mechanics of an infarcted ventricle.^58,59^ In our baseline EHT cocultures of cardiomyocytes and fibroblasts, the tissues exhibited significant contractile strains when electrically paced. After cryowounding a central portion of the tissues, we found the wounded regions began to stretch (positive tensile strain) while the remote zones continued to contract with each beat. In addition, the central wounded portion showed greater stiffness properties compared to the remote zones and the unwounded controls.

Of course, there are limitations to consider. One limitation is the use of immunofluorescence intensity as a surrogate metric for approximating cell survival after the cryo-wound. This approach lacks the specificity to distinguish between necrosis, apoptosis, viability, proliferation, etc. A more accurate approach would utilize specific cell death, proliferation, metabolic markers, thereby affording a more accurate display of the location, severity, and mechanism of cell turnover. Furthermore, this disease model is predicated on the use of a thermal injury. This method does not fully characterize the insults and the corresponding biochemical responses cells would experience under ischemic injury in vivo. But cryo-injury does provide a highly controllable model to induce a consistent wound size, shape, and position to achieve post-MI mechanical heterogeneity. Finally, this approach did not encompass direct evaluative methodologies to characterize the maturation of cardiomyocytes. Instead, it relies on alterations in force generation as an indirect measurement of cardiomyocyte development. While informative, this fails to capture the nuance of sarcomere alignment and underlying hypertrophy. Similarly, fibroblast activation was not directly measured in the form of fibroblast to myofibroblast transition.

Future directions for this platform should initially focus on deepening the characterization of both fibroblasts and cardiomyocytes in response to these different insults. A host of measurements to assay cellular cytoskeletal changes, gene expression changes, metabolic performance, morphology, etc. remains to demonstrate how closely the cells in our constructs match in vivo changes post-MI. Excitingly, the platform will enable these characterizations with human cells including iPSC-derived cardiomyocytes and fibroblasts derived either from human iPSCs or harvesting directly from patients. This is notable given the wide variability across animal models of MI.^10^ Ultimately, this 32-well in vitro bioreactor can be applied not just for testing basic scientific hypotheses but also for developing new therapeutic strategies as a drug-screening platform.

## 5. Conclusion

The platform we describe here utilizes a cryowound technique and regional analyses to investigate spatial heterogeneity in the response of cardiac fibroblasts and cardiomyocytes to infarct within a 3D hydrogel. These findings provide insight into the biomechanical changes that occur following myocardial infarction. The platform offers an opportunity to characterize the dynamic changes between cardiac cells in wound repair and to better understand the role of these cells in modulating the biomechanical properties of infarcted myocardium.

## 6. Acknowledgements

This work was performed at Clemson University within the Bioengineering Department. The authors wish to disclose inventor roles on patent application US-20230064704-A1 related to this work. This work was supported by funding from the NIH (R01HL144927 and P20GM121342), Clemson University Research Foundation, South Carolina Research Authority.

## Notes

### Competing Interest Statement

The authors wish to disclose inventor roles in patent application US-20230064704-A1 related to this work.

